# Global genetic patterns reveal host tropism versus cross-taxon transmission of bat Betacoronaviruses

**DOI:** 10.1101/2020.05.04.076281

**Authors:** Michael G. Bacus, Stephen Adrian H. Dayap, Nikki Vanesa T. Tampon, Marielle M. Udarbe, Roberto P. Puentespina, Sharon Yvette Angelina M. Villanueva, Aleyla E. de Cadiz, Marion John Michael M. Achondo, Lyre Anni E. Murao

**Affiliations:** Department of Biological Sciences and Environmental Studies, University of the Philippines Mindanao, Tugbok District, Davao City; Department of Medical Microbiology, College of Public Health, University of the Philippines Manila; Animal Solutions Veterinary Hospital, Bolcan, Davao City; Wildlife-Human Interaction Studies, Ecological Research and Biodiversity Conservation Laboratory, University of the Philippines Mindanao, Tugbok District, Davao City; Philippine Genome Center Mindanao, University of the Philippines Mindanao, Tugbok District, Davao City

**Keywords:** Coronaviruses, Bats, Phylogeny

## Abstract

Emerging infectious diseases due to coronavirus (CoV) infections have received significant global attention in the past decade and have been linked to bats as the original source. The diversity, distribution, and host associations of bat CoVs were investigated to assess their potential for zoonotic transmission. Phylogenetic, network, and principal coordinate analysis confirmed the classification of betacoronaviruses (BetaCoVs) into five groups (2A to 2E) and a potentially novel group, with further division of 2D into five subgroups. The genetic co-clustering of BetaCoVs among closely related bats reflects host taxon-specificity with each bat family as the host for a specific BetaCoV group, potentially a natural barrier against random transmission. The divergent pathway of BetaCoV and host evolution suggests that the viruses were introduced just prior to bat dispersal and speciation. As such, deviant patterns were observed such as for 2D-IV, wherein cross-taxon transmission due to overlap in bat habitats and geographic range among genetically divergent African bat hosts could have played a strong role on their shared CoV lineages. In fact, a few bat taxa especially the subfamily Pteropodinae were shown to host diverse groups of BetaCoVs. Therefore, ecological imbalances that disturb bat distribution may lead to loss of host specificity through cross-taxon transmission and multi-CoV infection. Hence, initiatives that minimize the destruction of wildlife habitats and limit wildlife-livestock-human interfaces are encouraged to help maintain the natural state of bat BetaCoVs in the wild.

**Importance:** Bat Betacoronaviruses (BetaCoVs) pose a significant threat to global public health and have been implicated in several epidemics such as the recent pandemic by severe acute respiratory syndrome coronavirus 2. Here, we show that bat BetaCoVs are predominantly host-specific, which could be a natural barrier against infection of other host types. However, a strong overlap in bat habitat and geographic range may facilitate viral transmission to unrelated hosts, and a few bat families have already been shown to host multi-CoV variants. We predict that continued disturbances on the ecological balance may eventually lead to loss of host specificity. When combined with enhanced wildlife-livestock-human interfaces, spillover to humans may be further facilitated. We should therefore start to define the ecological mechanisms surrounding zoonotic events. Global surveillance should be expanded and strengthened to assess the complete picture of bat coronavirus diversity and distribution and their potential to cause spillover infections.

## Introduction

Emerging and re-emerging infectious diseases greatly affect public health and global economies (1). These diseases involve pathogenic strains that recently evolved, pathogens that infect human population for the first time, and pathogens that re-occur at higher frequency (2). Majority of these emerging infectious diseases are caused by microorganisms from non-human source or zoonotic pathogens from wild animals (3). In particular, emerging infectious diseases due to coronavirus (CoV) infections have been receiving significant global attention as exemplified by the severe acute respiratory syndrome coronavirus (SARS-CoV) outbreak in 2002-2003, the Middle East respiratory syndrome coronavirus (MERS CoV) outbreak in 2012, and the recent SARS-CoV2 pandemic which causes the Coronavirus Disease of 2019 (COVID-19), all of which have been linked to bats as the original source (4-7).

Coronaviruses (CoVs) are pleiomorphic, single-stranded positive-sense RNA viruses with three major structural proteins; a nucleocapsid protein (N) which functions in encapsidating genomic RNA and facilitating its incorporation into virions, a small integral membrane protein (M) with intrinsic membrane-bending properties that plays a central role in viral assembly, an envelope glycoprotein (E), and a large spike protein (S) which functions in viral entry and pathogenesis (8-11). CoVs are considered to have the largest genome among RNA viruses at approximately 27 to 30 kb (6). There are four genera by which CoVs are classified, namely Alphacoronaviruses, Betacoronaviruses, Gammacoronaviruses and Deltacoronaviruses (12). Betacoronaviruses (BetaCoVs) are of particular importance as the SARS-CoV, MERS-CoV, and SARS-CoV2 which have caused global epidemics belong to this lineage (5). Betacoronaviruses (BetaCoVs) are further classified into subgenera Embecovirus (also lineage 2A and includes Murine CoV and ChRCoV HKU24), Sarbecovirus (lineage 2B and includes SARS-related CoVs), Merbecovirus (lineage 2C and includes Ty-BatCoV HKU4, Pi-BatCoV HKU5, Hp-BatCoV HKU25, and MERS-related CoVs), Nobecovirus (lineage 2D and includes Ro-BatCoV HKU9 and Ro-BatCoV GCCDC1), and Hibecovirus (lineage 2E and includes Bat Hp-betaCoV Zhejiang2013) (13).

HCoV-229E and HCoV-OC43, both human coronaviruses (HCoVs), were first discovered in patients with mild respiratory illness (14). Two new species of HCoVs, the HCoV-NL63 and HCoV-HKU1 were also discovered in 2004 and 2005, respectively (15). Disease types caused by HCoVs usually range from gastrointestinal infections, upper respiratory infections, and lower respiratory infections such as pneumonia (16). Further studies revealed that CoVs cause respiratory, enteric, hepatic and neurological diseases in animals like bats, birds, cats, dogs, pigs, mice, horses, and whales (17). Moreover, the SARS-CoV and MERS-CoV which have clear zoonotic origins have also been found to cause lower respiratory infections such as pneumonia. In particular, SARS-CoV has been associated with diffuse alveolar damage (DAD) and acute respiratory distress syndrome (ARDS), while MERS-CoV has been linked to renal failure (16). Recently, SARS-CoV2 has been reported to cause pneumonia, ARDS and multiple organ failure (7,18). CoVs can be transmitted via fecal-oral route, respiratory, as well as contact transmission (19). The spread of SARS-CoV has been primarily attributed to human-human transmission via direct contact with respiratory droplets and exposure to fomites (20). Similarly, although no evidence of being sustained, human-human transmission has also been reported for MERS-CoV infections (21). The World Health Organization (WHO) released guidelines on how to limit human-human transmission, and reduce the risk of animal-human transmission in order to contain the rapidly spreading COVID-19 disease that started in Wuhan, China (22).

There is increasing evidence for the role of bats as hosts of emerging pathogens, specifically viruses (23). Bats (Order: Chiroptera) are one of the most diverse and widely distributed animals, second only to rodents as the most speciose order in class Mammalia (24). They are classified into two suborders, Yinpterochiroptera, which include the false vampire bats (Megadermatidae), horseshoe bats (Rhinolophidae), and megabats or fruit bats (Pteropodidae); and Yangochiroptera which includes vesper bats (Vespertilionidae), sac-winged bats (Emballonuridae) and bulldog bats (Noctilionidae) (25). A significant number of CoVs can be found in bats, thus future spill-over events presents a constant threat to global health (26). In particular, emerging human coronaviruses have been linked to bat sources. Although camels were the source of the MERS-CoV in the Middle East, a coronavirus with 100% nt identity to that of human β-CoV 2c EMC/2012 isolated from a case-patient has been found in bats at a detection rate of 3.5% (27). SARS-like coronaviruses with 92% identity to that of human SARS-CoV isolates have also been detected in horseshoe bats in China. Three species of horseshoe bats from China namely *Rhinolophus pearsoni, R. pussilus* and *R. macrotis* demonstrated 28%, 33% and 78% SARS-CoV seroprevalence, respectively (28). Furthermore, SARS-CoV2 was found to be 96% identical to a bat coronavirus at the whole genome level (29). Bat CoVs (BtCoVs) comprise only 6% of the current CoV database although roughly 3,000 genetic lineages of BtCoVs are believed to circulate worldwide (30). It has also been suggested that future CoV outbreaks can be geographically predicted based on the specific bat species distribution (12). These highlight the need to expand our knowledge on BtCoVs, particularly on their diversity, distribution, host association, and evolution to understand their potential for zoonotic transmission. In this study, these parameters were assessed by classifying representative and unresolved BetaCoVs based on network and phylogenetic analysis of their RNA-dependent RNA polymerase sequences and evaluating patterns of geographic and host distribution. The analysis validated the current classification scheme of BetaCoVs with potentially novel groupings and subgroupings identified. Comparative phylogenetics demonstrated a strong tendency towards host specificity of bat BetaCoVs although there was poor evidence of co-evolution with their hosts.

## Results

### CoV detection in fruit bats from Southern Philippines

Small and large intestine samples were collected from 49 bat individuals, 67.35% of which belong to the lesser dog-faced fruit bat *Cynopterus brachyotis* mostly from residential sites but also present in the agricultural and forest sites, 20.41% to *Rousettus amplexicaudatus* all from agricultural sites, and 10.2% to the long-tongued nectar bat *Macroglossus minimus* mostly collected from agricultural sites (Table 1). Only one (2.04%) cave nectar bat *Eonycteris spelaea* was collected, which was captured in a forest site. Out of the 49 fruit bats tested, seven (14.29%) were positive for BtCoV based on RT-nPCR detection, all of which were from the bat species *C. brachyotis* (Table 1). The species-level detection rate of CoV among the *C. brachyotis* samples was 21.2% (7 out of 33), with five individuals positive for the small intestine samples, and two other individuals positive for the large intestine samples (Table 1). Most of the CoV-positive bats were females and juveniles that were captured in residential and forest sites near a watershed reservation (supplemental data).

**Table 1.**
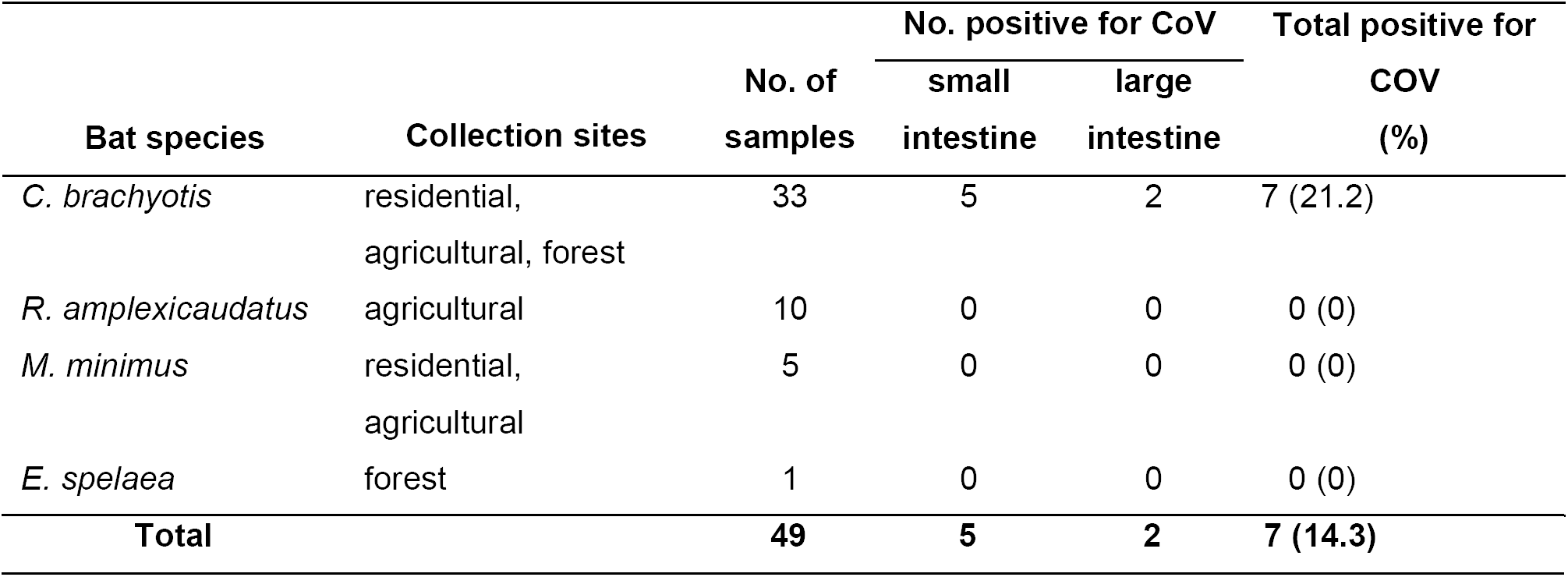
Summary of fruit bats collected from Malagos, Davao City and CoV detection.

### Phylogenetic relationships of global CoVs

A phylogenetic tree based on the partial RdRp gene sequences of CoVs obtained from this study and those mined from the NCBI database representing CoVs from various bat species, domestic and wild animals, as well as MERS-CoV, SARS-CoV and the SARS-CoV2, all from human patients was generated (Figure 1). Alpha and BetaCoVs each formed a distinct lineage from a common ancestor. Based on reference sequences, BetaCoVs diverged to five major phylogenetic clades classified as 2A (Embecovirus), 2B (Sarbecovirus), 2C (Merbecovirus), 2D (Nobecovirus), the recently proposed 2E (Hibecovirus) (13), and an unresolved clade which formed a genetic cluster distinct from the rest of the currently recognized BetaCoV subgroups. The same CoV groupings were supported by the network analysis using median-joining (Fig. 2A) and principal coordinate analysis using the distance matrix (Fig. 2B), wherein sequences from each clade and sub-clade formed corresponding unique and distinct clusters. Clade 2A was composed of human CoV (HCoV-OC43), porcine CoV (JL/2008), BtCoV (KX285045), and cattle CoV (NC_003045). Clade 2B was composed of human SARS-CoV, SARS-CoV2, SARS-like CoVs, and unclassified BtCoVs, while clade 2C was composed of human MERS-CoV, camel MERS-like CoVs, BtCoVs such as HKU5-1, and unclassified BtCoVs. BtCoVs such as HKU9 and GCCDC1 formed clade 2D along with many unclassified BtCoVs, which further diverged into five distinct subgroupings 2D-I to 2D-V. Clade 2E was composed of Bat HP-BetaCoV/Zhejiang2013 which is currently the only recognized strain that belongs to the subgenus Hibecovirus (13). Finally, the unresolved clade is composed of unclassified BtCoVs. The phylogenetic tree captured the current classification scheme for BetaCoVs with novel information on Nobecovirus subclassification. However, deviant samples were also observed such as BetaCoVs from *Rhinolophus pusillus* and *Myotis dasycneme* that clustered with the AlphaCoV lineage as consistently demonstrated by the phylogenetic, network, and principal coordinate analyses (Fig. 1 and 2).

**Figure 1.**
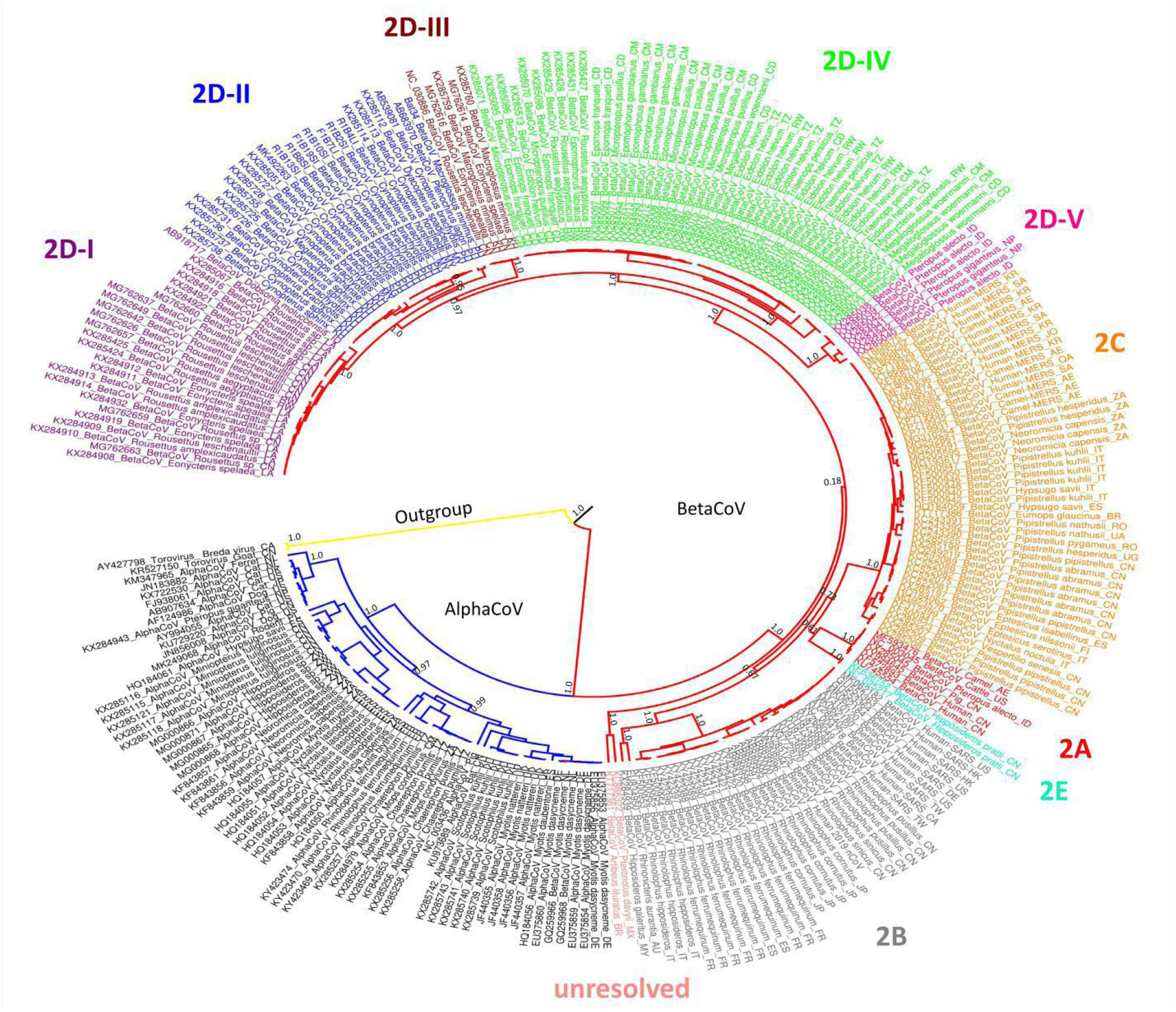
Bayesian phylogenetic tree of a 325 bp portion of the COV RdRp gene sequences available in GenBank and the samples obtained from this study. The betacoronavirus lineage is shown in red lines, while the alphacoronaviruses are shown in blue lines. The outgroup is represented by torovirus as shown in yellow line. Posterior values of the major divergence points are shown in the branches.

**Figure 2.**
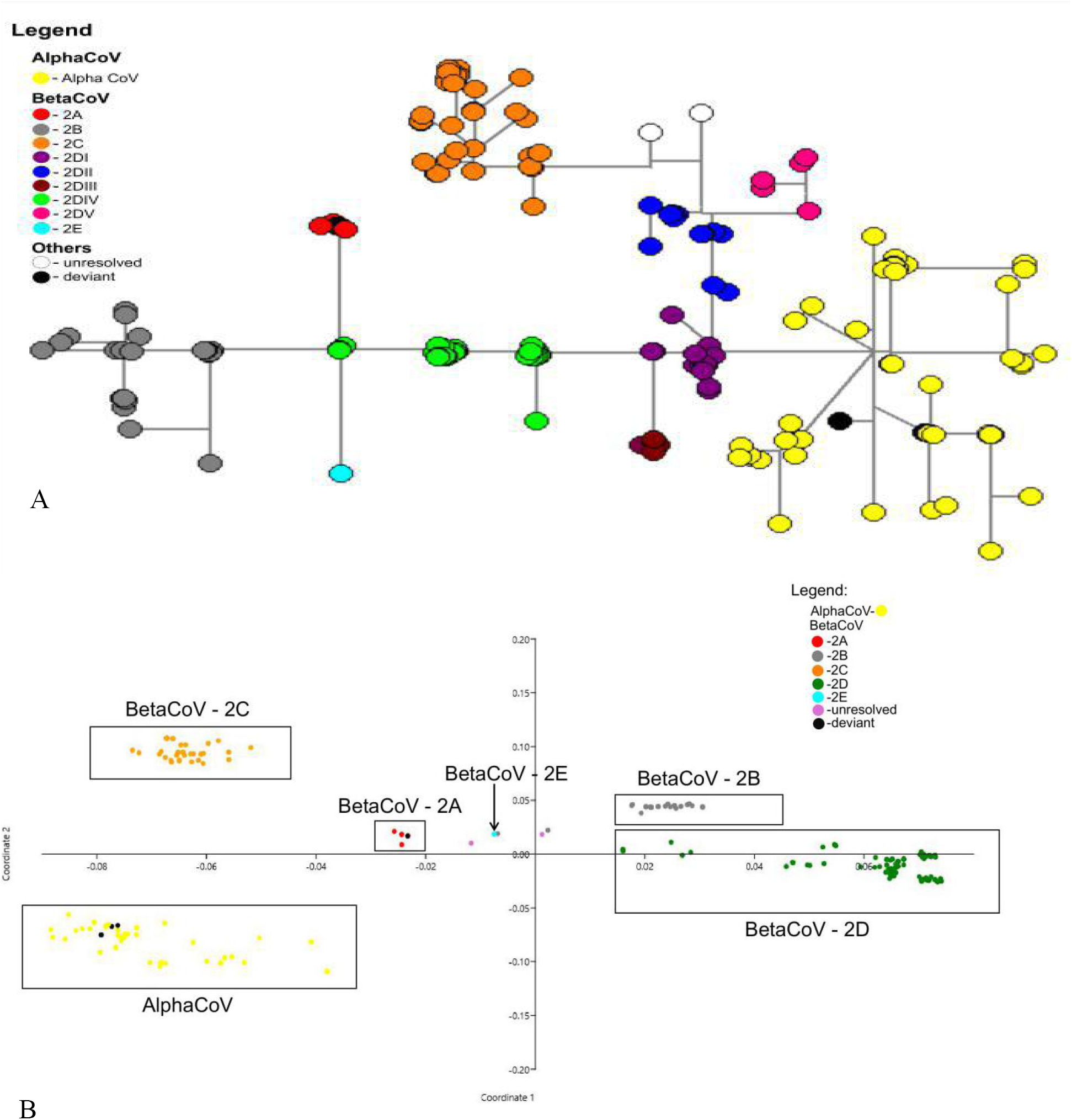
Genetic clustering of global CoVs using, a) network analysis through median joining of RdRp gene sequences, and b) principal coordinate analysis of the distance matrix of the RdRp gene sequences.

### Geographical distribution of bat CoVs

The regional distribution of bat CoVs comprising the major clades and subclades were examined using the network analysis as shown in Figure 3 and Table 2. Results showed a heterogeneous geographical distribution of CoVs for most of the clades, except for the 2D-II and 2D-IV subclade, which were exclusively found in bats from Southeast Asia and Africa, respectively. In contrast, Clade 2B or the Sarbecoviruses was distributed in Europe, East Asia, Southeast Asia, and Australia. Clade 2C or the Merbecoviruses also had regionally diverse distribution in East Asia, Middle East, Europe, Africa, and South America. The fruit bat subclade 2D-I also has a wide distribution from Africa, East Asia, and Southeast Asia, 2D-III in East and Southeast Asia, and 2D-V in South and Southeast Asia. The single representative CoV in 2E was from East Asia. The mammalian CoVs in 2A were from East Asia, Southeast Asia and North America, while the unresolved clade consisted of unclassified BtCoVs from South America.

**Table 2.**
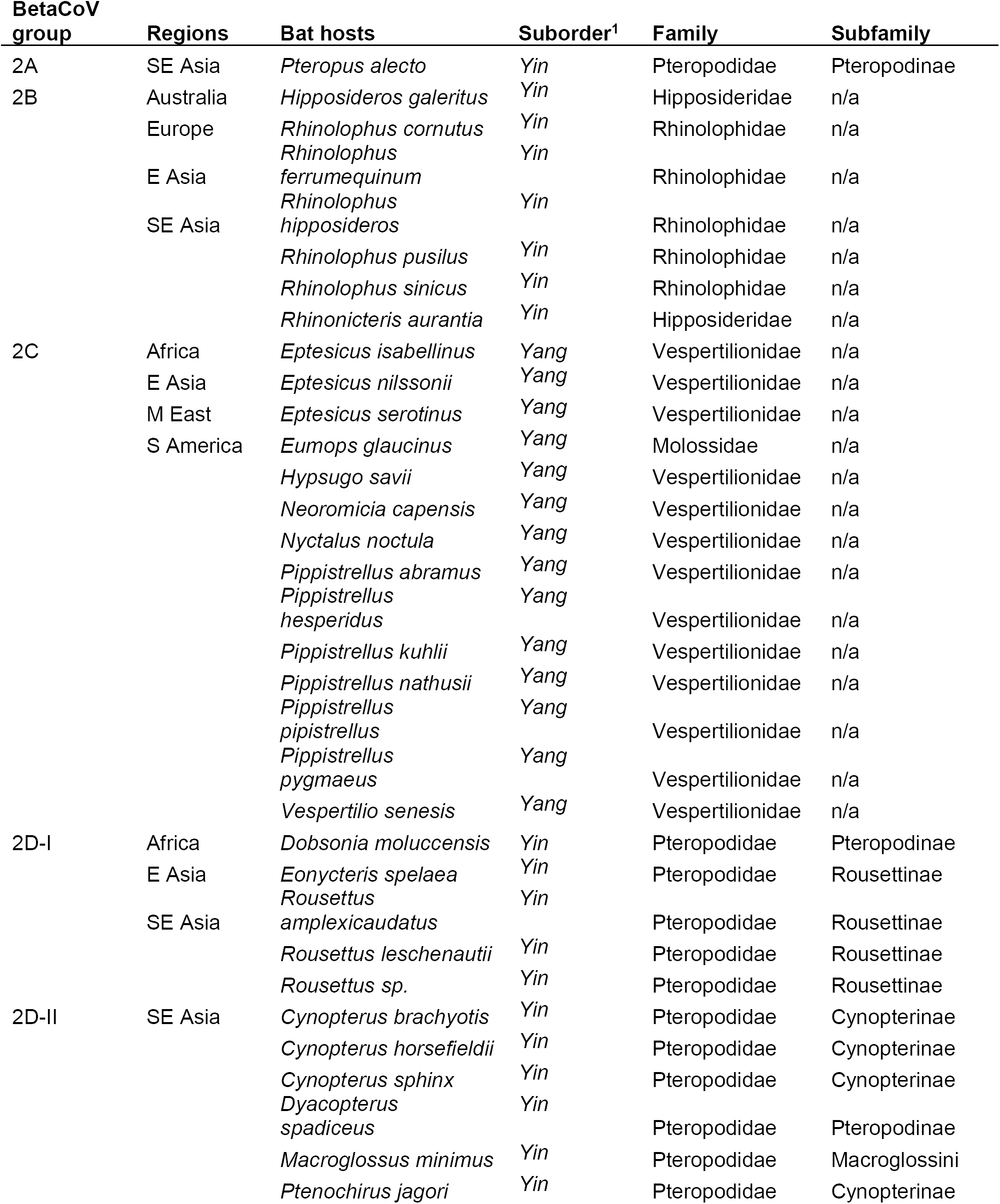
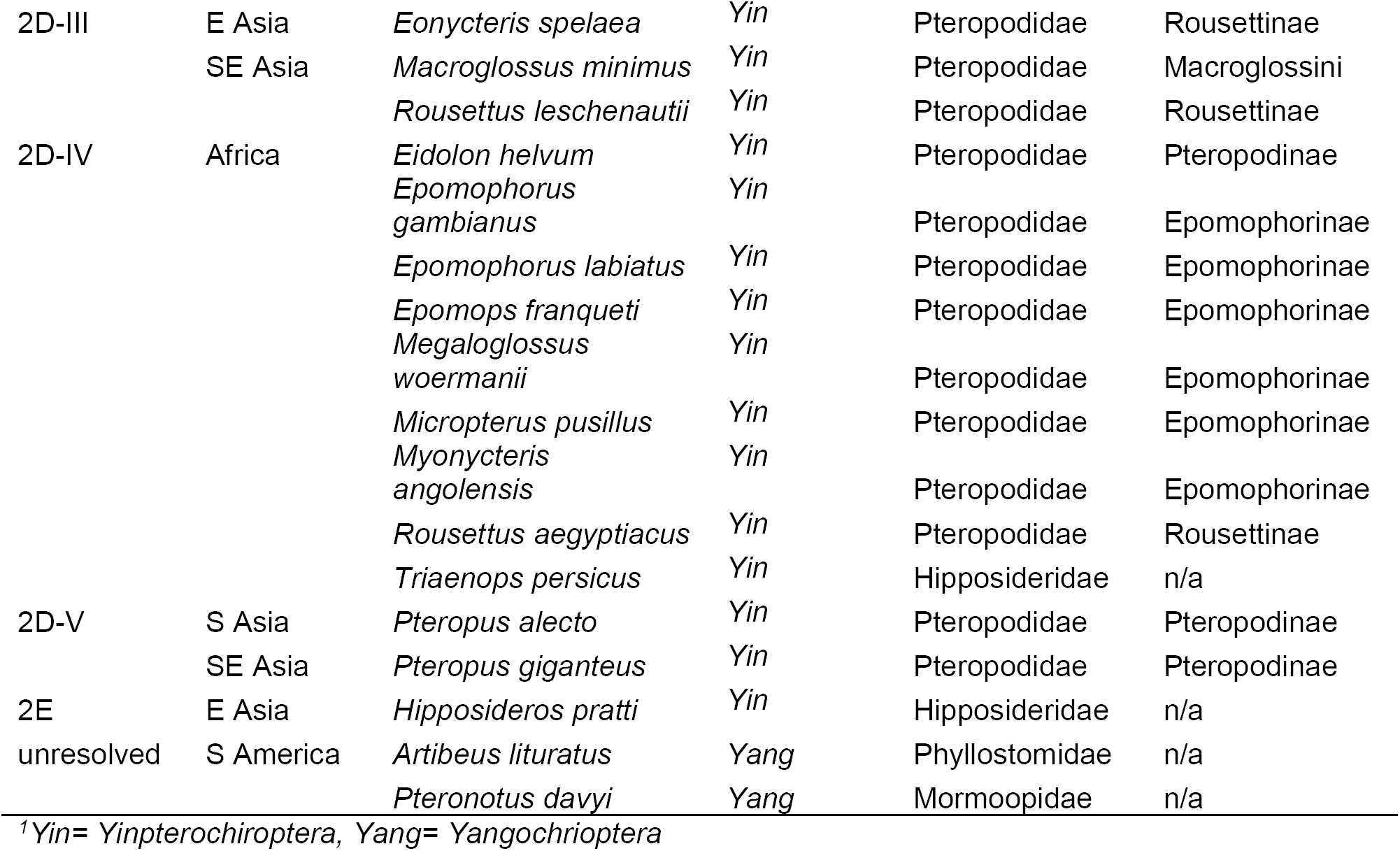
Summary of betacoronavirus groups and their corresponding regions and hosts

**Figure 3.**
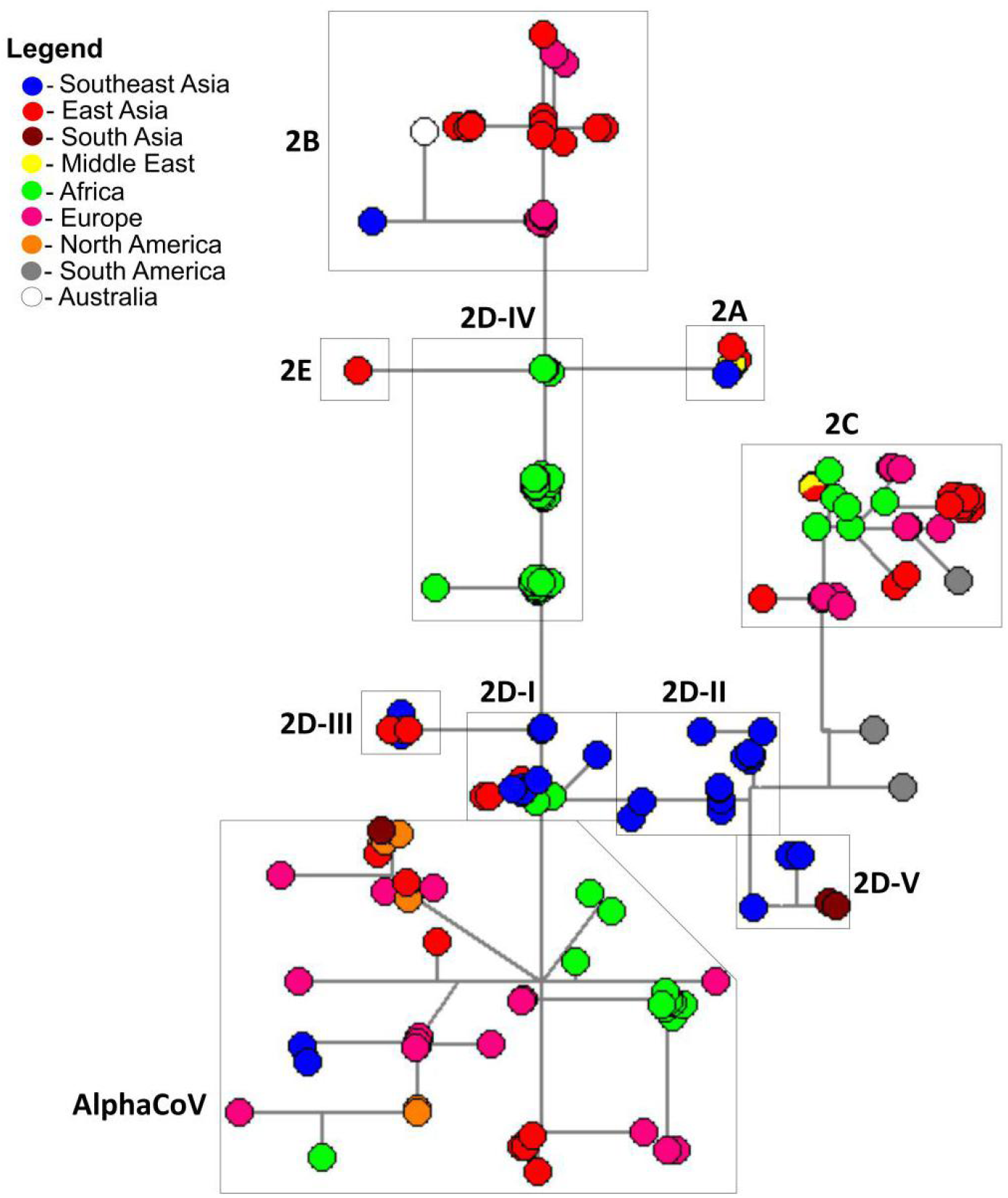
Regional distribution of CoVs from bats, domestic and wild animals, as well as humans using network analysis of RdRp gene sequences.

### Bat hosts of BetaCoV groups

The bat hosts were further evaluated to determine common patterns within the BetaCoV lineages. Network analysis revealed a heterogenous composition in most of the clades or subclades in terms of the bat source (Fig. 4 and Table 2). Clade 2A included one BtCoV from *Pteropus alecto*. Clade 2B, which includes the human SARS-CoV and SARS-CoV2, was primarily composed of BtCoVs from horseshoe bats (family Rhinolophidae) belonging to *Rhinolophus* sp. (67%), along with some Old World leaf-nosed bats (family Hipposideridae) such as *Rhinonicteris aurantia* and *Hipposideros galeritus*. Clade 2C of MERS-CoV was primarily composed of vesper bat hosts (family Vespertilionidae) such as *Pipistrellus* sp., *Neoromicia capensis, Hypsugo savii, Nyctalus noctula, Eptesicus* sp., and *Vespertilio sinensis,* wherein majority were sampled from *Pipistrellus* (46%). A CoV from *Eumops glaucinus* of the free-tailed bats (family Molossidae) was also found to cluster with this group. Meanwhile, clade 2D consisted primarily of fruit bat hosts (family Pteropodidae) and showed distinct subgroupings. CoVs from subfamilies Rousettinae (*Rousettus* sp. and *Eonycteris spelaea*), Pteropodinae (*Dobsonia* sp.), *a*nd Macroglossini (*Macroglossus minimus*) formed the subclade 2D-I and 2D-III, majority of which were sampled from the genus *Rousettus* (67%). CoVs from subfamilies Cynopterinae (*Cynopterus* sp., *Dyacopterus spadiceus, Megaerops niphanae,* and *Ptenochirus jagori*), *a*nd Macroglossini (*Macroglossus minimus*) formed subclade 2D-II, with the genus *Cynopterus* (83%) as the predominantly sampled group. 2D-IV was composed of CoVs mostly sampled from the African bat *Eidolon helvum* (subfamily Pteropodinae) (32%), and the rest from other African fruit bats of subfamily Epomophorinae (*Micropteropus pusillus, Epomophorus* sp., *Epomops franqueti, Megaloglossus woermanni,* and *Mynonycteris angolensis*), subfamily Rousettinae (*R. aegyptiacus*), and family Hipposidiridae (*Triaenops persicus*). Subclade 2D-V was composed solely of CoVs from *Pteropus* sp., commonly known as flying foxes (subfamily Pteropodinae), while the sole BtCoV representative in clade 2E has been detected in *Hipposideros pratti* (family Hipposidiridae). Finally, the unresolved clade was composed of American leafed (family Phyllostomidae) and mustached (family Mormoopidae) bat hosts.

**Figure 4.**
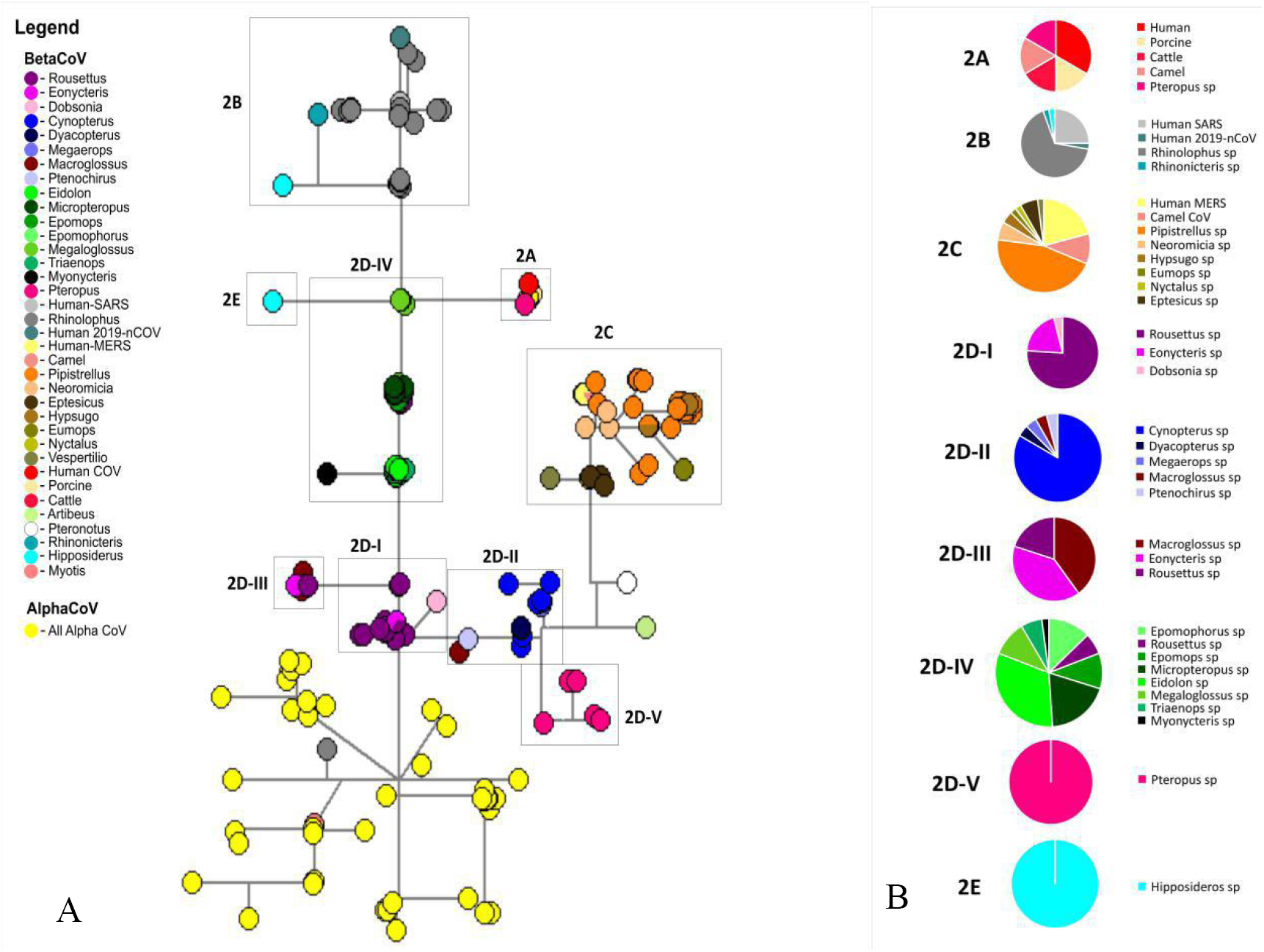
Host composition of betacoronavirus lineages as illustrated in the (a) network analysis of RdRp gene sequences, and (b) pie chart distribution of hosts per clade or subclade.

### Comparative phylogenetics of BetaCoVs and their bat hosts

Phylogenetic analysis using the *cytB* gene was subsequently conducted to understand the evolutionary relationships of the bat hosts within BetaCoV lineages (Fig. 5A). The microbats (suborder Yangochiroptera) formed two distinct lineages: the vesper bats versus the American leafed, mustached, and free-tailed bats. The megabats (suborder Yinpterochiroptera) also formed lineages corresponding to the three families: horseshoe, Old World leaf-nosed, and fruit bats. Furthermore, the fruit bats were sub-divided accordingly into Macroglossini, Pteropodinae, Cynopterinae, Rousettinae, and Epomophorinae lineages, except for *E. helvum* which was separated from the Pteropodinae group.

**Figure 5.**
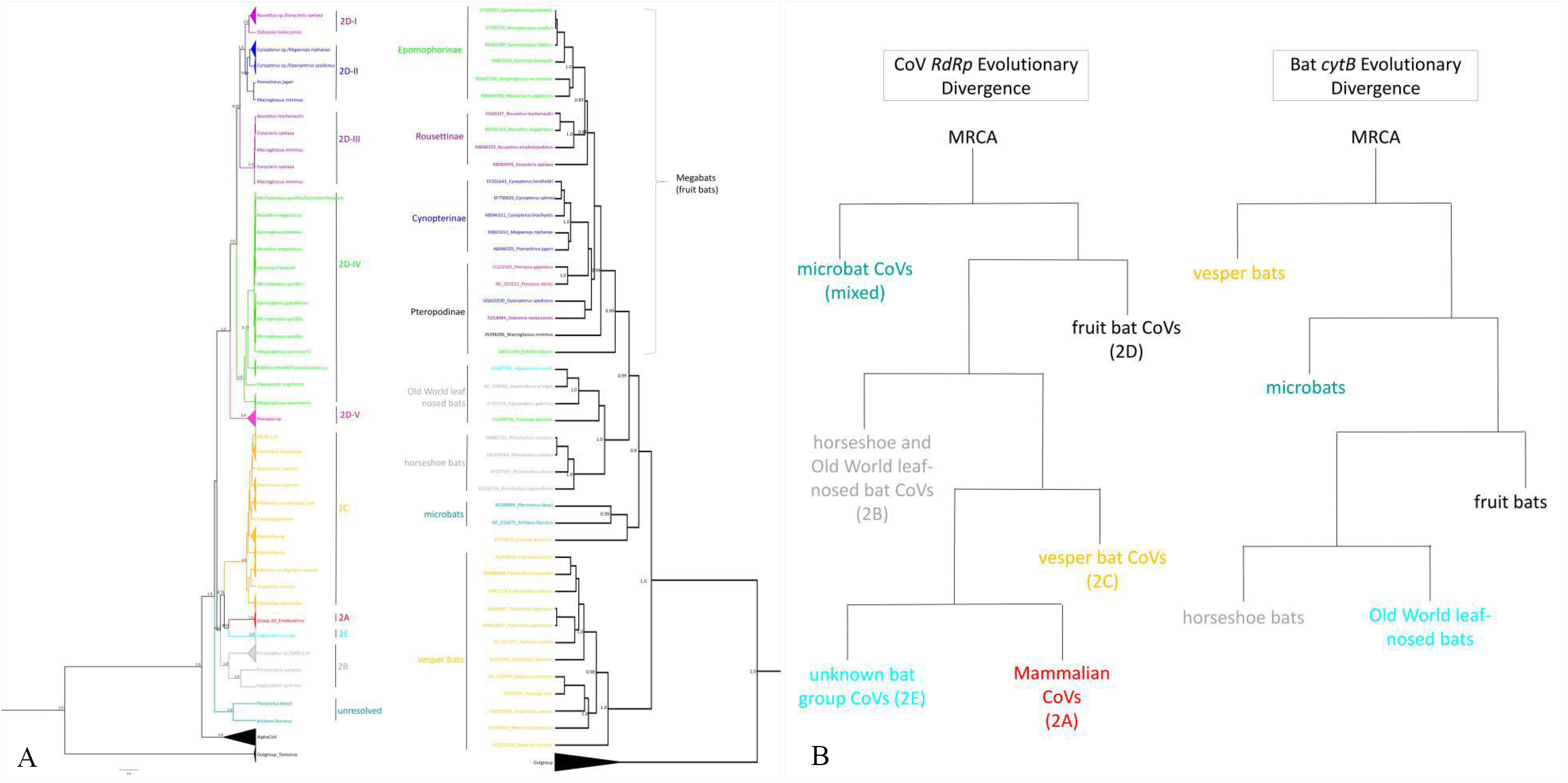
Phylogenetic relationships between CoVs (*RdRp* gene) and their bat host (*Cyt B* gene), A) bayesian phylogenetic trees generated using bayesian inference (BEAST v.1.10.4) with the GTR+G+I DNA substitution and site heterogeniety model, strict molecular clock and coalescent constant size, with posterior values of the well supported clades written in the nodes, B) evolutionary divergence patterns of CoVs and their bat hosts based from the generated Bayesian phylogenetic trees.

In general, there was a genetic clustering of bat hosts found within BetaCoV lineages. The 2B BetaCoVs was comprised of horseshoe and Old World leaf-nosed bat hosts that shared a common ancestor. A similar pattern was observed for 2C and the 2D subgroups. The 2C BetaCoVs was composed of vesper bat hosts, 2D-I/2D-III of Rousettinae bat hosts, 2D-II of Cynopterinae bat hosts, and 2D-IV of Epomophorinae bat hosts, wherein each bat group also formed their corresponding genetic clade. The 2D-V subgroup was composed of solely *Pteropus* sp. which belongs to Pteropodinae. Finally, the unresolved CoV clade was represented by American leafed and mustached microbat hosts (*P. davyii* and *A. lituratus*) that clustered with a common ancestor. Meanwhile, 2E was represented by only one bat host: *Hipposideros* sp. of the Old World leaf nosed bats.

However, some deviations were also noted. For example, bat hosts that belong to genetically unrelated taxa were mixed in some BetaCoV groups. The Mollosidae bat *Eumops glaucinus* was found in 2C BetaCoV of vesper bats, the Pteropodinae *Dobsonia moluccensis* in 2D-I of Rousettinae bats, the Pteropodinae *Macroglossus minimus* both in 2D-II of Cynopterinae bats and 2D-III of Rousettinae bats, and the Pteropodinae *Eidolon helvum*, Rousettinae *Rousettus aegyptiacus*, and Hipposideridae/Old World leaf-nosed bat *Trianeops persicus* in 2D-IV of Epomophorinae bats. Looking at the host, certain bat families were observed to harbor BetaCoVs that belong to various lineages. The Rousettinae bats were found to carry both 2D-I/2D-III and 2D-IV BetaCoVs, and the Old World fruit bats 2B, 2D-IV, and 2E. The subfamily Pteropodinae also hosted Nobecoviruses from various groups such as 2A (*P. alecto*), 2D-I (*D. moluccensis*), 2D-II (*D. spadiceus* and *M. minimus*), 2D-III (*M. minimus*), 2D-IV (*E. helvum*), and 2D-V (*Pteropus* sp.).

The divergence of the bats and their BetaCoVs were compared to evaluate common evolutionary pathways (Fig. 5B). The vesper microbats diverged as a separate group from the rest of the bats. The remaining bats further diverged into different clades: the first one comprised of the other microbat families (American leafed, mustached and free-tailed bats), and the second clade splitting into horseshoe bats, Old World leaf-nosed bats, and fruit bats, with the former two sharing a much recent common ancestor. For the corresponding viruses however, the evolutionary pattern was different. The unresolved microbat CoVs were the first to diverge from the rest of the bat BetaCoVs. The vesper microbat CoVs (2C) on the other hand appear to have diverged together with the Old World leaf-nosed megabat CoVs (2E). Finally, the mammalian CoVs (2A) emerged from a lineage of bat BetaCoVs.

## Discussion

A global analysis was conducted for BetaCoVs of human, animal, and bat origins including a complete set of representatives from the updated CoV classification (13). The major clades inferred from the generated Bayesian phylogenetic tree using partial RdRp sequences was consistent with previously reported classification of BetaCoVs (2A or Embecoviruses, 2B or Sarbecoviruses, 2C or Merbecoviruses, 2D or Nobecoviruses, and 2E or Hibecoviruses) using whole genome sequences of representative strains (12) and was able to position formerly unclassified bat CoVs. A new clade of unclassified bat BetaCoVs that is genetically distinct from the currently recognized groups was also observed. This classification can correct the GenBank annotation of deviant samples such as the BetaCoVs that grouped with AlphaCoVs, and add information on the currently unclassified BetaCoVs in GenBank. This is also the first report on the comprehensive classification of Nobecoviruses, of which there are currently only two recognized groups: Ro-BtCoV HKU9 and Ro-BtCoV GCCDC1 (12). Our analysis of unclassified BetaCoVs suggests that Nobecovirus diversity may have been underestimated in previous reports. We therefore propose the subclassification of Nobecoviruses into five subgroups (2D-I to 2D-V), which can help update surveillance records and facilitate monitoring of CoV populations in the wild.

The congruent association between the genetic clustering of bat CoVs and their bat hosts at different taxonomic levels (family, subfamily, and genus) and regardless of location suggests bat taxon-specific BetaCoV lineages: horseshoe and Old World leaf-nosed bats for 2B Sarbecoviruses, vesper bats for 2C Merbecoviruses, fruit bats for 2D Nobecoviruses, and the closely related American leafed and mustached bats for the new BetaCoV clade. Similar trends were observed upon further analysis of the Nobecoviruses and their subclades: 2D-I and 2D-III BetaCoVs were found mostly in Rousettinae bats, 2D-II in Cynopterinae bats, and 2D-V in *Pteropus* bats. A conclusive finding could not be generated from the single representative Old World leaf-nosed bat for 2E Hibecovirus. These findings support previous reports (31) which we have expanded here to identify the specific bat taxon associated with each BetaCoV group and a more extensive analysis of fruit bat CoVs. Network analysis showed no clear trends in the geographical distribution of closely related BetaCoVs except for 2D-IV, which is composed of four genetically distinct bat hosts (family Pteropodidae subfamily Pteropodinae, Epomophorinae, and Rousettinae; and family Hipposideridae), but all of which are found in Africa. Hence, host specificity could play a major role in BetaCoV diversity, except for 2D-IV for which geography may have a stronger influence.

Host specificity is uncommon in other bat-infecting viruses such as Paramyxoviruses and Papillomaviruses, which have been reported to have prevalent host switches (32,33). Indeed, bats have a predominant viral sharing network, suggesting that cross-species transmission events are common (34). In contrast, our findings support a previously proposed hypothesis that CoVs limit cross-species transmission within related bat taxa (31), which is indicative of a preferred adaptation to a certain range of hosts. Various lines of evidence have indicated bat-specific infectivity of CoVs. In one study, BtCoVs from primary infection of *C. brachyotis* had a reduced level of replication when experimentally inoculated to *Rousettus leschenaultii* (35). Similarly, the SARS-like WIV1-CoV, which was isolated from *Rhinolophus sinicus* bats and demonstrated positive replication in *R. sinicus* cell lines, showed weak infection in *Rousettus* sp. bats and cell lines (36-38). Indeed, our analysis showed that BetaCoVs from *Rousettus* sp. (2D-I, 2D-III or 2D-IV) are genetically distinct from *Cynopterus* sp. (2D-II) or *Rhinolophus* sp. (2B). Analysis of bat SARS-like CoV proteins demonstrated the absence of any selective pressure for evolution (39), suggesting that bat CoVs have already reached optimal fitness in their host. Factors that limit transmission to a different host taxon include cell surface receptors, the immune response, and replication fitness (40). These factors may serve as important natural barriers in the transmission of bat BetaCoVs, which can advantageously limit host jumping and co-infection, which otherwise may generate new virus strains with the ability to switch animal hosts (41,42).

Although bat BetaCoVs are host taxon-specific, their evolutionary pathways are different from bats, as has been reported in another study (31), suggesting that the virus did not have a long-term co-evolution with its host. Instead, this is indicative that the currently circulating viruses may have been introduced relatively recently, i.e. to the most recent common ancestors of each bat taxon but prior to global dispersion and speciation, during which the virus acquired adaptation to its host. The recent introduction of BetaCoVs in bats implies that other factors may have had the opportunity to influence virus-host dynamics. In the succeeding discussions, we will present two deviant phenomena that exemplify this: cross-taxon transmission of CoVs and bat hosts with multi-CoV lineages.

We provide genetic evidence for cross-taxon transmission as indicated by genetically unrelated bats that host BetaCoVs of the same lineage. For example, the Mollosidae bat *Eumops glaucinus* and vesper bats harbored CoVs that belong to 2C. *E. glaucinus* has a wide distribution in South America that overlaps with the vesper bat *Eptesicus sp.* (43,44), which is an opportunity for cross-species transmission of CoVs. Another example is the Pteropodinae bat *Dobsonia moluccensis, Dyacopterus spadiceus*, and *M. minimus* that carried Nobecoviruses from various subgroups. *Dobsonia sp.* and Rousettinae bats are primarily cave dwellers that have been documented to co-roost in the same cave habitat (45-49), which could explain their shared BetaCoVs from lineage 2D-I . Evidence of forage site overlap has also been documented for *Cynopterus sp.* and other Pteropodinae bats such as *M. minimus* which also feeds on banana and palm trees, and *Dyacopterus spadiceus* which have been documented to feed together with *Cynopterus horsfieldii* in the same foliage site (45). Here, we reported that these Cynopterinae and Pteropodinae bats have BetaCoVs that commonly belong to 2D-II. A final example is 2D-IV, which is composed of four genetically distinct bat hosts from Epomophorinae, Pteropodinae (*E. helvum*), Rousettinae (*R. aegyptiacus*), and Hipposidiridae/Old World leaf-nosed bats (*T. persicus*). The diverse hosts of 2D-IV could be explained by overlaps in foraging, roosting, and distribution of African bats. For example, *Rousettus aegyptiacus* has been reported to share the same foraging site with *Epomophorus gambianus* (50). Meanwhile, *Eidolon helvum* roosts gregariously, exhibiting an annual migration pattern as a function of food supply and has a wide distribution range in the African continent (51-53), which overlaps with the distribution and range of other Epomophorinae fruit bats in Africa. *Triaenops persicus* is also widely distributed in Eastern Africa, Middle East, and South Western Asia (54,55). Altogether, these imply that cross-taxon transmission of CoVs is most likely facilitated by close interactions brought about by an overlap of roosting and foraging habitat as well as geographic range of the bat hosts.

Our analysis also revealed that certain bat taxa such as Rousettinae, Hipposideridae (Old World fruit bats), and Pteropodinae harbor multi-CoV lineages. We hypothesize that certain host factors such as conserved cellular receptor motifs unique to some bat families are accessible to various forms of the CoV Spike protein (S) specifically the corresponding receptor-binding motif (RBM), thereby predisposing them to infection by various CoV lineages. It has been reported that different receptor-binding S1 subunit C-terminal domains (S1-CTDs) from different coronavirus lineages can recognize the same receptor (56). For example, the Lys353 amino acid in the angiotensin-converting enzyme 2 (ACE 2) plays a crucial role for the binding of the SARS-CoV and the HCoV-NL63, both of which are very divergent coronaviruses, the former being a Betacoronavirus, and the latter belonging to the alphacoronavirus lineage. Independent evolution of different RBDs in coronaviruses could lead to the recognition of the same virus-binding hotspot (56). Bats that can host a wide range of CoVs have the potential to propagate novel viruses. It is therefore recommended that the BtCoV database be expanded through sustained surveillance efforts covering more bat species especially from these three families in order to determine their full range of CoV lineages.

The phylogenetic analysis in this study points to bats as potential origins of other mammalian CoVs (2A). Switching to another taxon would require specific genetic alterations that will facilitate infection of a different host species (40). This entails a strong selection pressure, and all the more when switching to other animal hosts (35-38). A good demonstration of this is the evolution of human SARS-CoV which is believed to have occurred in a stepwise fashion, with the spike protein undergoing early selective pressure probably to mediate the switch from animal to human hosts, followed by the RdRp in the late stages to facilitate a more efficient replication in humans (42). Furthermore, the human MERS-CoV EMC/2012 was found to replicate in *Artibeus jamaicensis* bats, as well as in various cell lines from different bat families that have never been reported to host MERS-CoV strains (37,57). Viral strains with broad-spectrum tropism such as human SARS and MERS CoV are the result of an evolutionarily acquired ability that combined the use of new receptors, host immune evasion, and efficient replication in various host species (36,38,57,58). Anthropogenic activities such as climate changes affect the distribution of previously geographically restricted disease vectors (59). Continued ecological imbalances that alter bat distribution may eventually lead to loss of host specificity for bat BetaCoVs through cross-taxon transmission and adaptation of multiple CoV lineages. Diverse wildlife-livestock-human interfaces created by urbanization (60) could further increase the selection pressure resulting to spillover events in human populations. For example, SARS-CoV likely evolved to infect humans through a series of transmission events due to close or sustained contact between humans and animals in a wildlife market in China (61). Furthermore, the recent SARS-CoV2 outbreak in China has been reported to originate from a seafood market in Wuhan with exposure to wild animals (62). Considering all these factors, another novel human CoV outbreak originating from bats is imminent. These highlight the need to monitor and maintain the natural state of CoVs in the wild by strengthening routine surveillance of circulating CoVs, proper urban planning to minimize the destruction of wildlife habitats, and limiting wildlife-livestock-human interfaces such as by controlling wildlife consumption.

The detection rate of BtCoVs has been reported at a range of 2-30% in bats from various Asian countries, wherein our 14.29% detection rate in Southern Philippines is within range (63-67). Moreover, the 21.2% detection rate of CoVs in *Cynopterus brachyotis* from this study is lower compared to two separate studies in Northern Philippines at 37% and 39%, respectively (35,63), but higher compared to other Southeast Asian countries such as Thailand (11%) and Singapore (5.6%) (65,66). *C. brachyotis* is locally abundant and widely distributed throughout urbanized and secondary forests in both South and Southeast Asian regions, and have a high fruit species diversity in its diet (68). It is therefore recommended to explore the incidence of CoVs in the wild population of *C. brachyotis* in the Philippines, which may present a higher risk for future spillover infection in animal and/or human populations due to their presence in urban communities. On the other hand, the absence of CoV detection in other fruit bat species in this study does not rule out the possibility of these bats as reservoirs. This could have been due to sampling bias, i.e. non-*Cynopterus* species comprised only ∼30% of the captured bats, which highlights the need to strengthen surveillance efforts for BtCoVs in individual countries to estimate the true burden of viral diversity and distribution. Should there be a novel zoonotic CoV arising from fruit bats such as *C. brachyotis*, it is predicted to be genetically distinct from SARS-CoV, SARS-CoV2, or MERS-CoV. However, this novel virus may be just as virulent or highly contagious.

## Materials and Methods

### Bat sampling and tissue collection

BtCoV samples from Southern Philippines were obtained from an exploratory surveillance. Five sampling sites consisting of two agricultural sites, two residential sites and one forest site were selected for bat collection at Malagos, Davao City. Forty-nine individuals, all fruit bats, non-threatened and non-endemic, were collected through purposive sampling for two nights on November 16 and 17, 2018 and their morphometric measurements were recorded. Bat samples collected were identified using the Key to the Bats of the Philippines by Ingle and Heaney (1992) (69). Bat samples were anesthetized through an intraperitoneal injection of 0.1 ml tiletamine-zolazepam and euthanized via cardiac exsanguination to obtain small and large intestine samples. Prior to the conduct of the study, Wildlife Gratuitous Permit (WGP No. XI-2018-07) and IACUC approval (protocol no.: 2018-019) were secured from the Department of Environment and Natural Resources XI and the Institutional Animal Care and Use Committee of the University of the Philippines Manila, respectively.

### Coronavirus detection

Genomic RNA was extracted from small intestine and large intestine samples using the SV Total RNA Isolation kit (Promega, USA) according to the manufacturer’s instructions. All RNA extracts were subjected to reverse transcription polymerase chain reaction (RT-PCR) using the one-step RT-PCR kit (Qiagen, USA) and PanCoV F2 (5’-AAR TTY TAY GGH GGY TGG-3’) and PanCoV R1 (5’-GAR CAR AAT TCA TGH GGD CC-3’) primers (70). The RT-PCR mix was prepared as follows: 0.4 µl One-step RT-PCR enzyme mix, 0.4 µl 10 mM dNTP mix, 2 µl 5x One-step RT-PCR buffer, 0.2 µl of 10 µM each of PanCoV F2 and PanCoV R1 primers, 4.0 µl of RNase-Free water and 3 µl RNA extracts for a total of 10 µl per reaction. The cycling conditions were as follows: 30 minutes at 50°C, 15 minutes at 95°C, 40 cycles at 94°C for 40 seconds, 48°C for 40 seconds and 72°C for 1 minute. The final extension was at 72°C for 10 minutes. Nested-PCR was subsequently performed to amplify a 435 bp bat specific region of the RNA dependent RNA polymerase (RdRp) gene using in-house designed primers BatCoV F1 (5’-TGACAGAGCACTGCCCAA-3’) and BatCoV R1 (5’-TTGTAACAAACAACGCCATC-3’) (71), and the 2X Taq Master Mix (Vivantis, Subang Jaya, Malaysia). The nPCR mix was prepared as follows: 5 µl 2X Taq Master Mix, 0.4 µl of 10 µM each of BatCoV F1 and BatCoV R1 primers, 2 µl one-step RT-PCR product and 2.2 µl nuclease-free water for a total of 10 µl per reaction. The cycling conditions for nPCR were as follows: 2 minutes at 94°C, 35 cycles of 40 seconds at 94°C, 40 seconds at 48°C, 1 minute at 72°C, and a final extension for 10 minutes at 72°C. The expected 435 bp amplicon of the BtCoV RdRp gene was visualized through electrophoresis using a 1.5% agarose gel.

### Sequence processing

Positive amplicons with an expected size of 435 bp were excised and purified using the GF-1 AmbiClean Kit (Vivantis, Subang Jaya, Malaysia) and were sent to Macrogen, Korea for standard DNA sequencing. Sequences were cleaned using the FinchTV software (Geospiza, USA) and distance analysis was performed using the Basic Local Alignment Search Tool (BLAST) of the National Center for Biotechnology Information (NCBI) (http://www.ncbi.nlm.nih.gov/).

### Phylogenetic analysis of global CoV sequences

CoV sequences obtained from this study, including 223 RdRp sequences of BtCoVs, 22 Human CoV sequences which include SARS, MERS and the 2019-nCoV isolated from patients, and 19 CoV sequences from other animal hosts such as camels, cats, dogs and pigs that were obtained from the National Center for Biotechnology Information (NCBI) database were used for phylogenetic analysis. Multiple sequence alignment was performed with the MAFFT software using the default algorithm and the leave gappy regions function. The final alignment was trimmed and cleaned in MEGA 7 software in order to come up with a 325 bp alignment of the CoV partial RdRp gene. The best phylogenetic model was calculated using jModelTest v.2.1.10 (72). The executable xml file for phylogenetic analysis was prepared using BEAUTi v.1.10.4 with the GTR+G+I DNA substitution and site heterogeneity model, which had the lowest Bayesian Information Criterion (BIC) as previously calculated in jModelTest, a length of chain of 100 million, strict molecular clock, and coalescent constant size model. The rest of the tree priors were set to the default value. Phylogenetic inference was performed using BEAST v.1.10.4 and the log files were evaluated using Tracer v.1.7.1 to see if the estimated sampling size (ESS) values for most of the continuous parameters are sufficient (>200) (73,74). A Maximum Clade Credibility (MCC) tree was generated using TreeAnnotator v.1.10.4. and the resulting MCC tree was visualized with FigTree v.1.4.4 (http://tree.bio.ed.ac.uk/software/figtree). Results from phylogenetic analysis were compared with principal coordinate analysis using the pairwise distance matrix of the CoV sequences using Past 3 (75) and with the network analysis using median joining in terms of major clades, geographical region and host genus.

### Phylogenetic analysis of bats

*Cytochrome B* (*Cyt B*) gene of 43 different bat species representing the bat hosts of the betacoronavirus lineage were obtained from the NCBI database. Phylogenetic analysis was performed in the same manner as previously described for CoV sequences using a GTR+G+I DNA substitution and heterogeneity model, which had the lowest Bayesian Information Criterion (BIC) as calculated in jModelTest. The resulting bat phylogenetic trees were plotted against the CoV phylogenetic tree to assess virus and host evolutionary congruence.

## Acknowledgement

This research was funded by the Department of Science and Technology - Philippine Council for Health Research and Development through the Regional Health Research and Development Council XI Grant No. 18-1572. The authors would like to thank the Local Government Units of Malagos Davao City, the Malagos Garden, the Philippine Eagle Foundation, Mr. Kemuel Libre Jr., Yancy Yurong and Alex Tiongco for all the assistance and support in during the field sampling of this study.

